# Dentate gyrus neurons that are born at the peak of development, but not before or after, die in adulthood

**DOI:** 10.1101/650309

**Authors:** Tina Ciric, Shaina P. Cahill, Jason S. Snyder

## Abstract

In the dentate gyrus of the rodent hippocampus, neurogenesis begins prenatally and continues to the end of life. Adult-born neurons often die in the first few weeks after mitosis, but then survive indefinitely. In contrast, neurons born at the peak of development are initially stable but can die later in adulthood. Physiological and pathological changes in hippocampal structure may therefore result from both the addition of new neurons and the loss of older neurons. However, it is unknown whether neurons born at other stages of development also undergo delayed cell death. Here, we used BrdU to label dentate granule cells that were born in rats on embryonic day 19 (E19; before the developmental peak), postnatal day 6 (P6; peak) and P21 (after the peak). We quantified BrdU^+^ neurons in separate groups of rats at 2 and 6 months post-BrdU injection. Consistent with previous work, there was a 15% loss of P6-born neurons between 2 and 6 months of age. In contrast, E19- or P21-born neurons were stable throughout young adulthood. Delayed death of P6-born neurons suggests they may play a unique role in hippocampal plasticity and pathology in adulthood.

---

Proposed functions of dentate gyrus neurogenesis have been inspired by patterns of cell addition and loss. In particular, neuronal survival has been repeatedly linked to behavior and cognitive demand. At baseline, a large proportion of adult-born neurons die within the first few weeks after mitosis (Cameron et al., 1993; Kempermann et al., 2003). Learning and environmental stimulation can rescue adult-born neurons at specific stages of cellular development, typically around 1-2 weeks of cell age (Gould et al., 1999; Epp et al., 2007; Tashiro et al., 2007; Anderson et al., 2010). Learning also causes death of specific cohorts of immature neurons, suggesting that both the addition and removal of neurons is involved in the adaptive response to experience (Olariu et al., 2005; Dupret et al., 2007). Survival of immature DG neurons is competitive and depends on NMDA receptors (Tashiro et al., 2006). Since mature adult-born neurons selectively respond to stimuli that they encountered at immaturity (Kee et al., 2007), they may store information as long as they survive, which appears to be indefinite (Dayer et al., 2003; Kempermann et al., 2003).

In contrast to our detailed understanding of the dynamics of neurogenesis in adulthood, much less is known about neurons born in early development, even though this is when the majority of DG neurons are born (Snyder, 2019). Specifically, while there have been many studies of DG development, few studies have examined the properties of age-defined cohorts of developmentally-born DG neurons. We recently examined the short- and long-term survival of rat DG neurons born at the peak of development, P6 (Cahill et al., 2017). Whereas adult-born neurons have an early critical period for survival, P6-born neurons did not. Also, while adult-born neurons are stable after reaching 4 weeks of age, we found that 17% of P6-born neurons died between 2-6 months of age. The functional relevance of this delayed cell death is unknown, but it could contribute to forgetting and hippocampal/DG atrophy in disorders such as depression (McKinnon et al., 2009; Boldrini et al., 2013; Mahar et al., 2017). Whether delayed cell death is observed among cells born at ages other than P6 is unknown.

Mature DG neurons are not identical and even cells born at different stages of perinatal development can have distinct properties (Kerloch et al., 2018; Save et al., 2018; Snyder, 2019). Indeed, DG neurogenesis spans many developmental milestones that may differentially recruit new neurons and affect their survival phenotype (e.g. birth, eye opening, weaning). To determine the scope of delayed cell death we therefore examined long-term survival of DG neurons born at three different stages of rat development: 1) DG granule neurons born on embryonic day 19. These are largely generated by proliferative cells that are migrating from the primary to secondary germinal zones, outside of the dentate gyrus (Altman and Bayer, 1990a). They are among the earliest granule neurons added to the DG. 2) DG granule neurons born on postnatal day 6. These cells are born in the tertiary germinal zone, which is located in the dentate hilus (Altman and Bayer, 1990b). Postnatal day 6 is within the the peak of DG development so cells born at this time represent a large proportion of DG neurons (Schlessinger et al., 1975). This population is known to undergo cell death in young adulthood (Dayer et al., 2003; Cahill et al., 2017). 3) DG granule neurons born at P21. These neurons are generated in the subgranular zone, at the border between the granule cell layer and the hilus (Altman and Bayer, 1990b). Adult neurogenesis also occurs in the subgranular zone. However, rats at P21 are sexually immature, still acquiring hippocampal learning and memory abilities (Akers and Hamilton, 2007; Raineki et al., 2010), and are therefore fundamentally distinct from adults.

Except where stated otherwise, methodological details are identical to those of our recent study of cell death in P6-born cells (Cahill et al., 2017). To label cells born prenatally (E19 group), rats were generated from timed pregnancies. Male and female breeders were paired each afternoon and females were inspected for the presence of sperm the following morning. The day when sperm was detected was designated as E1 and at this time females were separated from the males and housed singly for the remainder of the pregnancy. On E19, mothers were injected once with the thymidine analog BrdU (50 mg/kg, subcutaneous; Roche) to label proliferative cells that are beginning to migrate to the DG. Mothers remained with the litters for three weeks after birth, when offspring were weaned to 2 per cage. For the P6 and P21 groups, breeder pairs remained together from conception to age P21, when offspring were weaned 2/cage. For postnatal groups, the day of birth was designated as postnatal day 1. Pups were injected with BrdU on postnatal day 6 (50 mg/kg, intraperitoneal) or postnatal day 21 (200 mg/kg, intraperitoneal). Rats in the P21 group were injected as they were weaned into their new cages. Groups of rats from each condition (E19, P6, P21) were perfused at 8 weeks and 6 months post-BrdU injection. Thus, cell ages were identical across studies but animal ages varied slightly. We refer to the 8 week group as “2 months” for simplicity. Hippocampi were processed for BrdU immunohistochemistry as previously described (Cahill et al., 2017).

Total BrdU^+^ cell counts were estimated from coded slides with the optical fractionator technique and stereological counting principles (West et al., 1991), using an Olympus BX53 light microscope equipped with a motorized stage and software for stereological sampling (StereoInvestigator; MicroBrightField). Dentate gyrus granule cell layer areas were outlined at 2x magnification and cell counting was performed at 40x magnification. The section sampling frequency for all groups was 1/12. Other sampling parameters were optimized to accommodate differences in cell density across groups. Specifically, for the E19 group the sampling grid was 380 × 380 μm, the counting frame was 70 × 70 μm and the dissector height was 13 μm with a 3 μm guard zone. For the P6 group the sampling grid was 220 × 220 μm, the counting frame was 70 × 70 μm and the dissector height was 8 μm with a 4 μm guard zone. For the P21 group the sampling grid was 100 × 100 μm, the counting frame was 60 × 60 μm and the dissector height was 13 μm with a 3 μm guard zone.

BrdU injections resulted age-related patterns of cell labelling that were consistent with previous studies of rodent development (Fig. 1). Injections at E19 labelled CA3 and CA1 pyramidal neurons and DG neurons in the superficial granule cell layer of the dorsal DG. In the ventral DG, BrdU+ cells were located in deeper portions of the granule cell layer consistent with generation of granule neurons beginning at ~E14 and proceeding along ventral to dorsal, and superficial to deep, axes (Schlessinger et al., 1975; Altman and Bayer, 1990a). E19-born neurons were either darkly or lightly stained for BrdU. Darkly labelled cells most likely reflect cells that were undergoing mitosis at the time of BrdU injection and lightly labelled cells are either, a) those that were generated by subsequent divisions and underwent label dilution, or b) those that took up BrdU for only a portion of S-phase. P6 BrdU injections did not label any pyramidal neurons in the CA fields, as expected. In the DG, BrdU^+^ granule cells were primarily located in the middle ~50% of the granule cell layer, but labelled cells could be observed in the more superficial and deep aspects as well. P6-born cells had less variation in in BrdU staining intensity than E19 cells and most cells tended to be strongly and evenly stained throughout the nucleus. Finally, BrdU injections on P21 labelled a smaller population of cells that were primarily located in the deep portion of the granule cell layer. P21-born cells were typically located in the deep 1/3 of the granule cell layer, with some cells located between them and the hilus due to continued addition of younger cohorts of granule cells after P21.

**Figure 1:**
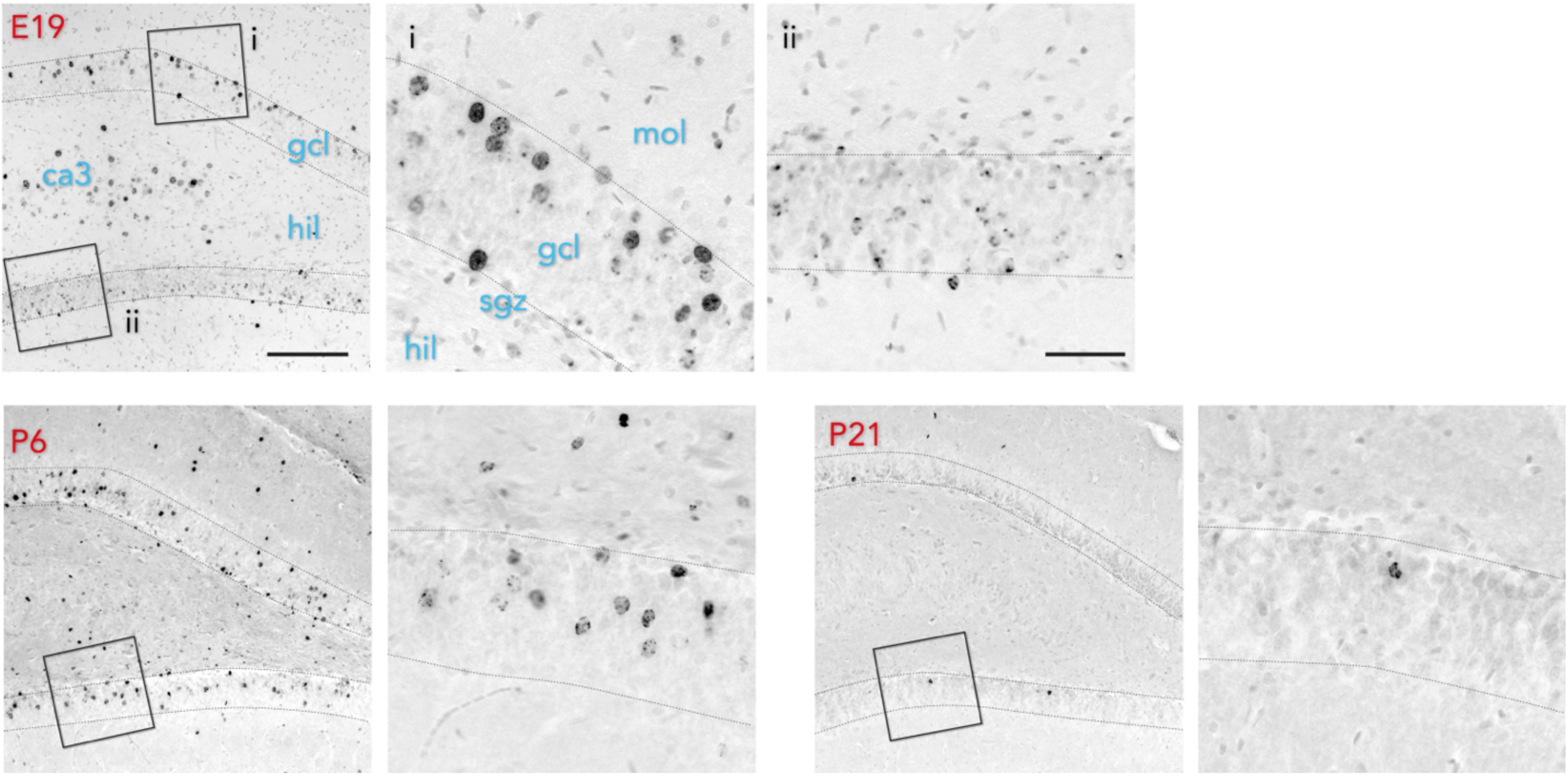
BrdU labelling of developmentally-born cells. E19 injections labelled granule neurons in the superficial granule cell layer, near the molecular layer. Strongly labelled cells can be seen in inset i. Weakly-labelled, speckled cells can be seen in inset ii. P6 injections labelled cells primarily in the middle of the granule cell layer. P21 injections labelled many fewer cells that were located in the deep portion of the granule cell layer, near the hilus/subgranular zone. gcl, granule cell layer; hil, hilus; mol, molecular layer; sgz subgranular zone. Scale bars 200 µm (low magnification images) and 50 µm (high magnification images).

Since E19 BrdU injections resulted in darkly and lightly labelled cells we set staining intensity criteria to avoid counting cells that were labelled due to repeated division of stem cells after E19. First, we only counted cells that had ≥ 10% of their nucleus stained for BrdU. This revealed 163,188 cells at 2 months and 153,929 cells at 6 months (coefficient of error = 0.06; T_15_=0.5, P=0.6). Since these numbers were comparable to our previous counts of P6-born cells (Cahill et al., 2017), and neurogenesis rates are ~3x lower at E19 than at P6 (Schlessinger et al., 1975), this criterion likely included many cells that were labelled by redivision and overestimated the true value. We therefore adjusted our criterion to include only cells that were unambiguously, strongly labelled with BrdU (≥ 50% of the nucleus stained). Stereological quantification of intensely labelled cells revealed ~45,000 granule neurons, which is likely a more accurate estimate of the true E19 value. While the coefficient of error increased when we only included the intensely-labelled cells (0.11), since fewer cells were sampled, counts were virtually identical at the 2 and 6 month time points. These results collectively indicate that cells born prenatally do not undergo delayed cell death in young adulthood (Fig. 2). Consistent with evidence that ~3x more cells are generated on P6 compared to E19 (Schlessinger et al., 1975), ~150,000 granule neurons were born on P6. There was a 15% loss of BrdU^+^ cells between 2 and 6 months of age, replicating our previous finding (Cahill et al., 2017). By P21, ~10,000 cells were labelled, which was over 10x lower than P6 levels. There was no loss of P21-born cells from 2 and 6 months of age. Error coefficients were low for both the P6 (0.06) and P21 (0.07) groups.

**Figure 2:**
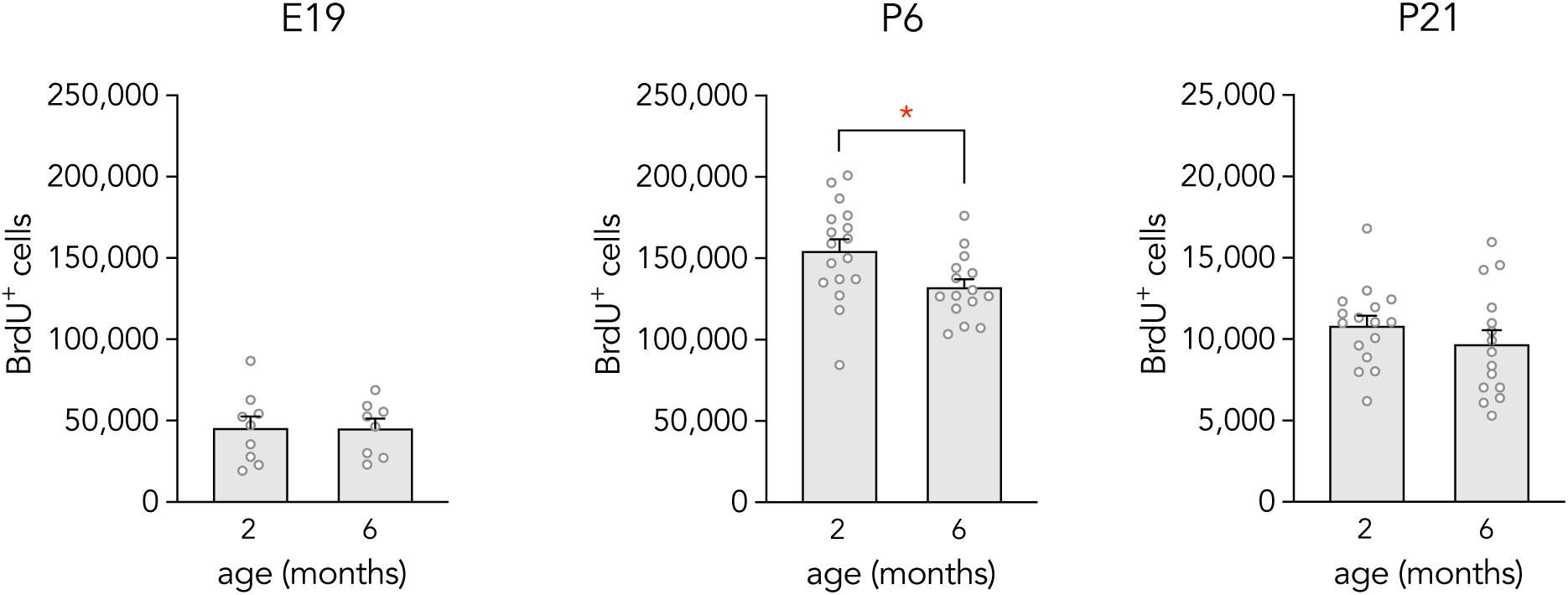
Long-term neuronal survival. A similar number of E19-born cells were present at 2 and 6 months of age (T_15_= 0.008, P= 0.99). There was a 15% loss of P6-born cells between 2 and 6 months of age (T_30_= 2.5, *P=0.02). A similar number of P21-born cells were present at 2 and 6 months of age (T_29_= 1.1, P=0.3). Note the 10x smaller y-axis scale for the P21 data. Bars represent mean ± standard error.

These results replicate previous findings that P6-born granule neurons undergo delayed cell death in young adulthood and they reveal that this cellular attrition is specific to neurons born around the peak of DG development. While our additional timepoints identify E19 and P21 as boundaries, the precise age window during which these “vulnerable” cells are generated, and therefore the total number that are lost, remains unclear. Only one study has quantified rat DG neurogenesis throughout the early developmental period (Schlessinger et al., 1975), finding that 1-3% of the granule cell population is generated daily between E14 and E21. Neurogenesis spikes on the day of birth and 5-8% of granule neurons are added daily between P1-P8 (7% on average). If similar patterns are observed for all cells born in this window, 8% of all granule cells would be lost in young adulthood, or ~160,000 cells. Of course, the window extends to older ages, even more cells may be lost. Notably, these rates of cell loss are broadly comparable to the rates of cell addition over the same timeframe, due to adult neurogenesis (345,000 cells added between 2-6 months (Snyder and Cameron, 2012)). Whether there is a causal balance between neuronal loss and addition remains to be investigated.

An outstanding question is why P6-born cells undergo delayed cell death but not cells born earlier or later in development. Assuming that there is a degree of functional heterogeneity within the DG that depends on when a cell was born (Snyder, 2019), P6-born cells may be reflective of the largest cohort of DG neurons, and therefore may be the most dispensable. Possibly, E19-born cells reflect specialized dentate gyrus cell populations (Kerloch et al., 2018; Save et al., 2018), which render them more resistant to cell death. It is also likely that E19-generated neurons are very different from the majority of DG neurons since they undergo significant migration to reach their final destination. Finally, a recent report indicates that P21-born neurons in mice are structurally identical to adult-born neurons (Kerloch et al., 2018), suggesting that, by weaning, neurogenesis produces a consistent granule cell phenotype.

Ultimately, functional studies of age-defined cohorts of neurons will be needed to determine the mechanisms and consequences of granule cell loss. The cells examined in this study are clearly developing in very different sensory environments. E19-born cells may be those cells that are developing afferent synapses during the 2^nd^ postnatal week, before eye opening (Ye et al., 2000). P6-born cells display peak immediate-early gene expression at P21, when animals are leaving the nest (Cahill et al., 2017). Finally, P21-born cells will likely be in their critical period of development and plasticity during adolescence and early adulthood. Early life experiences have lasting effects on hippocampal function (Nguyen et al., 2015). Thus, the retention and removal of cells that were developing during early life experiences may shape adaptive behavioral responses in adulthood.

## Acknowledgements

This research was supported by a Discovery Grant from the Natural Sciences and Engineering Research Council of Canada (NSERC; JSS), a New Investigator Award from the Canadian Institutes of Health Research (JSS), a Scholar award from the Michael Smith Foundation for Health Research (JSS) and a NSERC Postgraduate Scholarship and Killam Doctoral Scholarship (SPC).

## Data Availability

The data that support the findings of this study are available from the corresponding author upon request.

